# Targeting neuroinflammation by pharmacologic down-regulation of inflammatory pathways is neuroprotective in protein misfolding disorders

**DOI:** 10.1101/2022.09.26.509513

**Authors:** Sydney J. Risen, Sean W. Boland, Sadhana Sharma, Grace M. Weisman, Payton M. Shirley, Amanda S. Latham, Arielle J.D. Hay, Vincenzo S. Gilberto, Amelia D. Hines, Stephen Brindley, Jared M. Brown, Stephanie McGrath, Anushree Chatterjee, Prashant Nagpal, Julie A. Moreno

## Abstract

Neuroinflammation plays a crucial role in the development of neurodegenerative protein misfolding disorders. This category of progressive diseases includes, but is not limited to, Alzheimer’s disease, Parkinson’s disease, and prion diseases. Shared pathogenesis involves the accumulation of misfolded proteins, chronic neuroinflammation, and synaptic dysfunction, ultimately leading to irreversible neuronal loss, measurable cognitive deficits, and death. Presently, there are little to no effective treatments to halt the advancement of neurodegenerative diseases. We hypothesized directly targeting neuroinflammation by downregulating the transcription factor, NF-κB and the inflammasome protein, NLRP3, with the brain-penetrant, non-toxic, SB_NI_112, would be neuroprotective. To achieve this, we used a cocktail of RNA targeting therapeutics (SB_NI_112) shown to be brain-penetrant, non-toxic, and targeting both NF-|B and NLRP3. We utilized a mouse-adapted prion strain as a model for neurodegenerative diseases to assess aggregation of misfolded proteins, glial inflammation, neuronal loss, cognitive deficits, and lifespan. Prion-diseased mice were treated either intraperitoneally or intranasally with SB_NI_112. Behavioral and cognitive deficits were significantly protected by this combination of NF-κB and NLRP3 down-regulators. Treatment reduced glial inflammation, protected against neuronal loss, prevented spongiotic change, rescued cognitive deficits, and significantly lengthened lifespan of prion-diseased mice. We have identified a non-toxic, systemic pharmacologic that down-regulates NF-|B and NLRP3, prevents neuronal death and slows the progression of neurodegenerative disease. Though mouse models do not always predict human patient success, and the study was limited due to sample size and number of dosing methods utilized, these findings serve as a proof of principle for continued translation of the therapeutic SB_NI_112 for prion disease and other neurodegenerative diseases. Based on success in a murine prion model, we will be continual testing SB_NI_112 in a variety of neurodegenerative disease models, including Alzheimer’s Disease and Parkinson’s Disease.

## Introduction

Chronic inflammation in the brain has emerged as a significant contributor to neurodegenerative disease pathogenesis and progression. Alzheimer’s disease (AD), Parkinson’s disease (PD), and prion diseases share many common characteristics; accumulation and aggregation of misfolded proteins, glial activation, chronic neuroinflammation, neuronal loss, cognitive deficits, and death^1–5^. These diseases can be placed under the broader category of neurodegenerative protein misfolding disorders (neurodegenerative diseases) and are associated with the presence and aggregation of misfolded amyloid-β (Aβ) in AD, alpha synuclein (α-Syn) in PD, or the prion protein (PrP^Sc^) in the brain, which lead to behavioral deficits and subsequent progressive neuronal death^6^. Many disease modifying treatments have been directly targeted toward the aggregated proteins associated with such disease, though none have been successful in stopping disease progression^7–9^. Innate immune activation of microglia and astrocytes appears to be particularly important in the context of neurodegenerative diseases^10–12^. Furthermore, new evidence suggests that the compounded effect of misfolded proteins leads to further glial and astrocyte activation^6,13–15^. Therefore, chronic neuroinflammation has been identified as a promising therapeutic target for neurodegenerative diseases. However, the only current interventions include anti-inflammatory steroids, but their effectiveness as neurotherapeutic countermeasures has not yet been established physiologicaly^16,17^.

During neurodegenerative disease-induced brain stress, there is significant crosstalk between microglia, astrocytes, and neurons via critical inflammatory cell signaling events involving initiation of the nuclear factor kappa B (NF-κB) pathway and subsequent NOD (nucleotide-binding oligomerization domain), LRR (leucine rich repeat), and pyrin-containing 3 (NLRP3) inflammasome formation^18–20^. NF-κB, when translocated into the nucleus, promotes transcription of several inflammatory cytokines and chemokines, including NLRP3, pro-interleukin-1β (pro-IL-1β), and pro-interleukin-18 (pro-IL-18), triggering a cascade of microglial and astrocytic inflammation^21^. These reactive astrocytes are directly toxic to neurons and endothelial cells during prion disease^22,23^. Many recent studies, including human patient, *in vitro*, and *in vivo* animal models, have pointed to the assembly of the NLRP3 inflammasome in microglia and astrocytes as a significant contributor to inflammatory-related amplification of misfolded proteins in neurodegenerative diseases^24–31^. Formation of the NLRP3 inflammasome is a critical step for activation of pro-inflammatory cytokines and perpetuation of the immune response through microglial pyroptosis^32,33^. Pro-caspase-1 is activated by proximity-induced autocatalytic activation upon recruitment to the NLRP3 inflammasome. Active caspase-1 cleaves pro-IL-1β and pro-IL-18 into their mature and biologically active forms, cytokines IL-1β and IL-18^32,34^. In mouse models of APP/PS1 mutation and a tauopathy, NLRP3 inflammasome assembly has been identified as a significant contributor to inflammatory-related amplification of Aβ plaques and neurofibrillary tangles^24,25^. Similarly, the NLRP3 inflammasome promotes α-Syn production in mouse models of PD^26,27^. Activation of NF-κB in prion disease augments neuronal death by increasing the expression of pro-apoptotic genes and triggering the release of cytochrome C, resulting in mitochondrial dysfunction and endoplasmic reticulum stress^35^. Additionally, the upregulation of inflammatory cytokines IL-1β and tumor necrosis factor (TNF), formerly known as tumor necrosis factor alpha (TNFα), at sites of synaptic damage trigger the production of reactive oxygen species including hydrogen peroxide and nitric oxide, leading to excitotoxicity and neuronal death^36–38^. Beyond induction of inflammation and propagation of misfolded proteins, it has been proposed that activation of the NLRP3 inflammasome during NMPDs inhibits microglial autophagy, dampening the identification and degradation of misfolded protein aggregates in AD, PD, and prion disease^39–41^.

Based on the extensive evidence supporting central involvement of NLRP3 inflammasome formation following NF-κB activation in the progression of neurodegenerative diseases, it is important to develop potential therapeutics to inhibit NF-κB and NLRP3. Therapeutic intervention to target and inhibit NLRP3 inflammasome formation arises as a potential three-hit treatment against neurodegenerative diseases by: 1) reducing inflammatory signaling amongst microglia and astrocytes^18–20^, 2) preventing propagation of misfolded proteins^24–31^, and 3) upregulating microglia autophagy to accelerate removal of misfolded proteins^39–41^. No studies to date have explored combined NF-|B and NLRP3 inhibition as a therapeutic intervention for neurodegenerative diseases.

In the present study, we utilized non-toxic, brain-penetrant Nanoligomers designed to downregulate NF-κB and NLRP3 by blocking their translation^42^ (Figure 1A). Nanoligomers are stable, non-toxic, and bioavailable, with the capability to cross the blood brain barrier (BBB) through various routes of administration (Figure 1A)^43^. A previous study displayed effective gene regulation *in vivo* with a similar Nanoligomer therapeutic aimed at down-regulation of NF-κB and tumor necrosis factor receptor 1 (TNFR1) in an LPS-induced murine model of neuroinflammation^42^. The combined NF-κB and NLRP3 down-regulator, SB_NI_112, displays high target specificity and minimal off-targeting^42^. To study the impact of targeting NLRP3 inflammasome formation as a therapeutic for neurodegenerative diseases, we utilized a mouse model of infectious prion disease in wild-type (C57BL/6) mice. A prion model allows for study of a naturally occurring neurodegenerative diseases without the need for transgenic mice. Prion diseases are rare neurodegenerative diseases that result from the native conformation of the cellular prion protein (PrP^C^) misfolding to the infectious form, PrP^Sc^ ^44,45^. All prion diseases display the common characteristics of neurodegenerative diseases, including the accumulation and aggregation of a misfolded protein (PrP^Sc^), glial activation, chronic inflammation, neuronal loss, cognitive deficits, and death^42,43^. Previous literature has shown that therapeutics developed for prion disease mice can be translated into other neurodegenerative diseases, though the few compounds shown to reduce signs of prion disease in mouse models are associated with several negative side effects^36^. In this study, found treatment with SB_NI_112 to down-regulate NF-κB and NLRP3 reduced glial inflammation, protected against neuronal loss, prevented spongiotic change, rescued cognitive deficits, and significantly lengthened lifespan of prion-diseased mice. These findings, if validated clinically, have important implications for improving the quality of life for patients suffering with neurodegenerative diseases.

**Figure 1.**
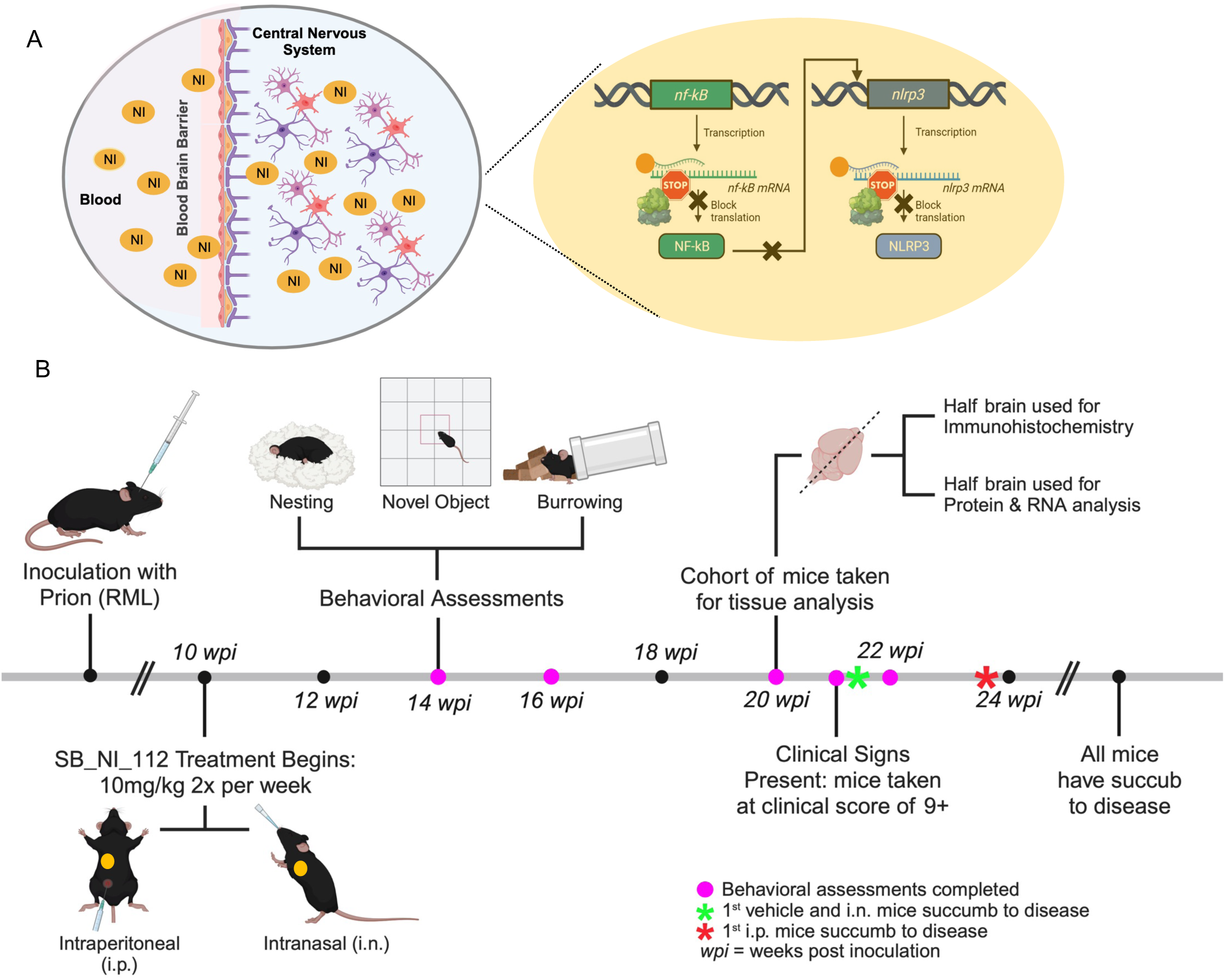
Experimental Design. **A)** Schematic demonstrating the ability of Nanoligomers to cross the BBB, as well as the mechanism of action (translational inhibition) through which Nanoligomers repress NFKB and NLRP3 expression. **B)** Wild-type C57BL/6 mice were inoculated with RML mouse-adapted prion or normal brain homogenate (NBH). Incubation period is 10 weeks. Prion diseased mice were treated with NF-|B and NLRP3 down-regulator SB_NI_112 or vehicle via intraperitoneal (i.p.) or intranasal (i.n.) routes of exposure, twice a week at 10 mg/kg starting at 10 weeks post inoculation (wpi). Mice were monitored for changes in cognition and behavior throughout the disease progression. At 20 wpi brain tissue was dissected for analysis of disease progression including glial inflammation, spongiotic change, and neuronal loss. Created with BioRender.com

## Results & Discussion

Attempts to slow the progression of neurodegenerative diseases have proven to be difficult. Many promising treatments are toxic or must be delivered directly to the brain, making translation to human patients difficult^7,36,46,47^. Neurodegenerative diseases have highly complex etiology, making the root cause, or multiple causes, difficult to identify and target. Neuroinflammation has arisen as a promising target, as chronic stimulation of the brain’s immune system leads to increased protein misfolding and neuronal death^6,11,15^. The NF-κB pathway and subsequent NLRP3 inflammasome formation have been identified as key contributors to chronic neuroinflammation present in neurodegenerative diseases^30,31,35,37^. Here, we employed a mouse prion model to test the efficacy of a combined NF-κB and NLRP3 down-regulator for neurodegenerative disease treatment.

Wild-type, C57Bl6/J, (Jackson Laboratories) mice were infected intracerebrally with 0.1% prion brain homogenates from Rocky Mountain Laboratories (RML) or normal brain homogenates (NBH) derived from C57Bl6/J mice as the controls. SB_NI_112 treatment began at 10 weeks post-inoculation (wpi) or approximately 40% of prion disease progression and continued until the mice succumbed to the disease (Figure 1B). As the disease progressed and treatment continued, behavioral assessments including novel object recognition, burrowing, and nesting, were performed at 12 wpi to train and establish a baseline for each experimental cohort, then continued throughout the study. All three behavioral assessments measure hippocampal integrity, the brain region known to show the first clinical pathology including glial inflammation and spongiotic change^48,49^.

### SB_NI_112 penetrates the blood-brain barrier and is systemically non-toxic

With a focus on reducing inflammation in the brain, it was important to assess whether SB_NI_112 effectively crosses the blood brain barrier (BBB). Biodistribution and toxicity testing was completed in naïve C57BL/6 female and male mice. Penetration of SB_NI_112 across the blood-brain barrier was confirmed with ICP-MS and detected in the hippocampus (76.1 nM +/- 44.3), cortex (49.9 nM +/- 11.4), cerebellum, (96.9 nM +/- 45.0) and brainstem (96.6 nM +/- 32.4) (Figure 2A), which is significantly higher than the dissociation constant of SB_NI_112^50^.

**Figure 2.**
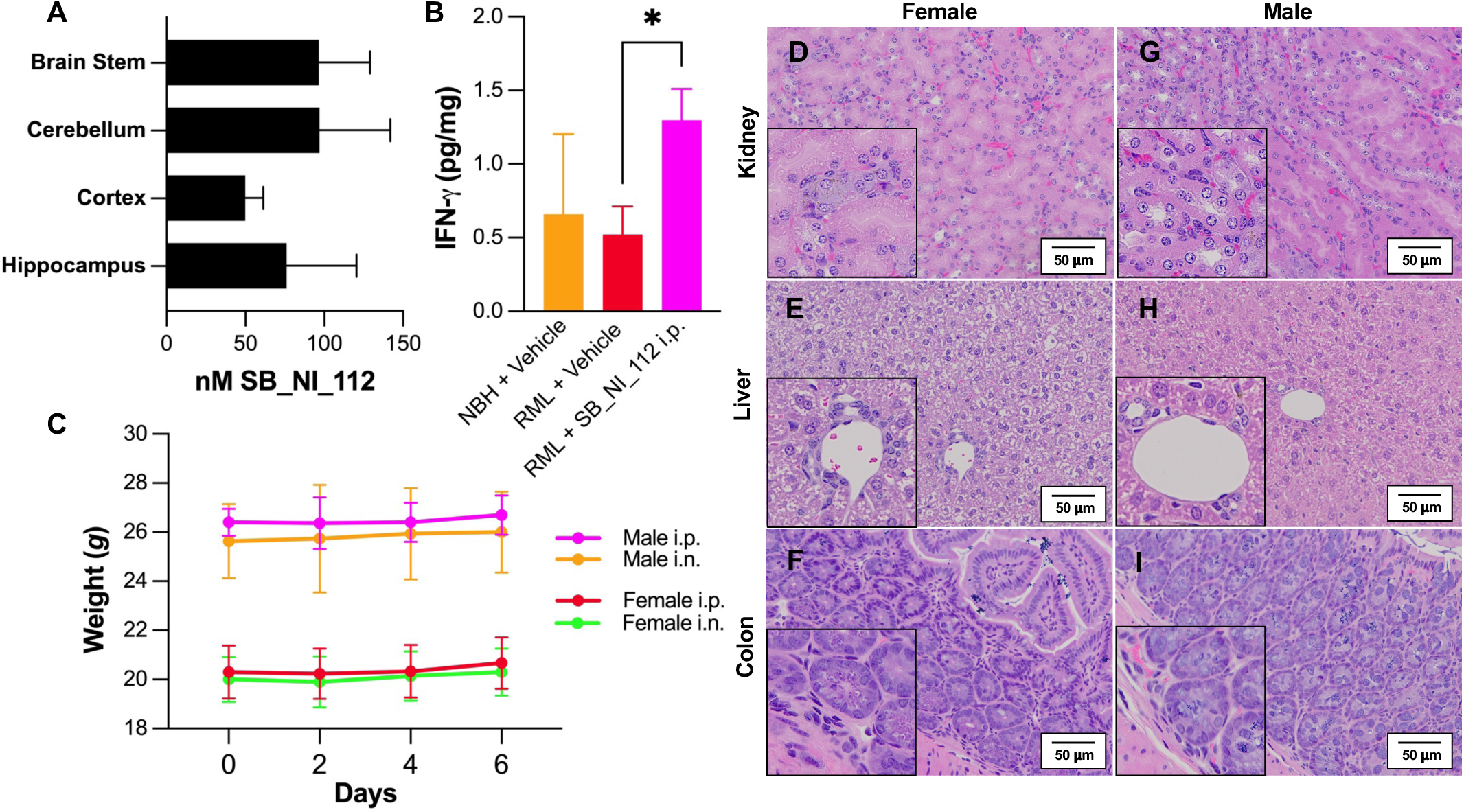
NF-|B and NLRP3 down-regulator SB_NI_112 penetrates the brain and is systemically non-toxic. Levels of SB_NI_112 in brain regions following dosing at 150 mg/kg i.p. three times per week for two weeks (A). Expression of IFN-© measured with ELISA (B). N=2 for NBH + Vehicle, N=4 for RML + Vehicle and RML + SB_NI_112 i.p. 10 mg/kg. One-way ANOVA, error bars = SEM, *p < 0.05. Weights of mice following a single dose of 500 mg/kg SB_NI_112 i.p. (C). Representative images at 20x of colon, kidney, and liver for female (D-F) and male (G-I) C57BL/6 mice one week after 500 mg/kg dose.

Down-regulation of NF-|B and NLRP3 with SB_NI_112 leads to dampening of chronic inflammation while preserving key immune pathways needed to respond to infectious pathogens (63). Despite down-regulation of NF-|B and NLRP3, interferon gamma (IFN-γ) expression was significantly higher in prion infected mice treated with SB_NI_112 (p = 0.048 (2-way t-test, Excel) or 0.0468 (One-way ANOVA, Graphpad Prism)), and not significantly different from control NBH mice (Figure 2B). This shows the use of SB_NI_112 specifically interferes with the NF-κB and NLRP3 pathway but does not alter expression of immune signaling molecules essential for response to infections.

Both female and male naïve C57BL/6 mice treated with a single dose of SB_NI_112 at 500 mg/kg intraperitoneally (i.p.) had no weight loss 1 week after dosing (Figure 2C) and displayed no morphologic toxicity in the colon (Figure 2C, F), kidney (Figure 2D, G), or liver (Figure 2E, H) via hematoxylin and eosin staining. No immune cell infiltration was identified in these organs and the parenchymal cells were intact. Based on previous studies completed with similar therapeutics^42,43^, we chose to dose mice conservatively at 10 mg/kg two times per week. Prion-diseased mice were split into 3 treatment groups: vehicle i.p., SB_NI_112, intranasal (i.n.), and SB_NI_112 i.p.

To confirm therapeutic SB_NI_112 successfully inhibits both NF-κB and NLRP3, we performed western blots to analyze the ratio of active NF-|B (phosphorylated p65; p-p65) compared to inactive form (p65) and NLRP3 protein expression. Interestingly, we identified a higher success of inhibition with intraperitoneal delivery than intranasal. SB_NI_112 provided significant reduction of p-p65 in prion disease mice compared to untreated counterparts (Figure 3A, B, and D: p-value = 0.0333, Figure S3). Intraperitoneal treatment also significantly reduced NLRP3 expression in prion-diseased mice compared to untreated counterparts (Figure 3C, E; p-value = 0.0380, Figure S3). This difference in effectiveness is likely due to inconsistency of intranasal delivery compared to the intraperitoneal method. The higher success of SB_NI_112 dosed intraperitoneally to inhibit NF-|B and NLRP3 is supported by the following behavioral and histological results obtained.

**Figure 3.**
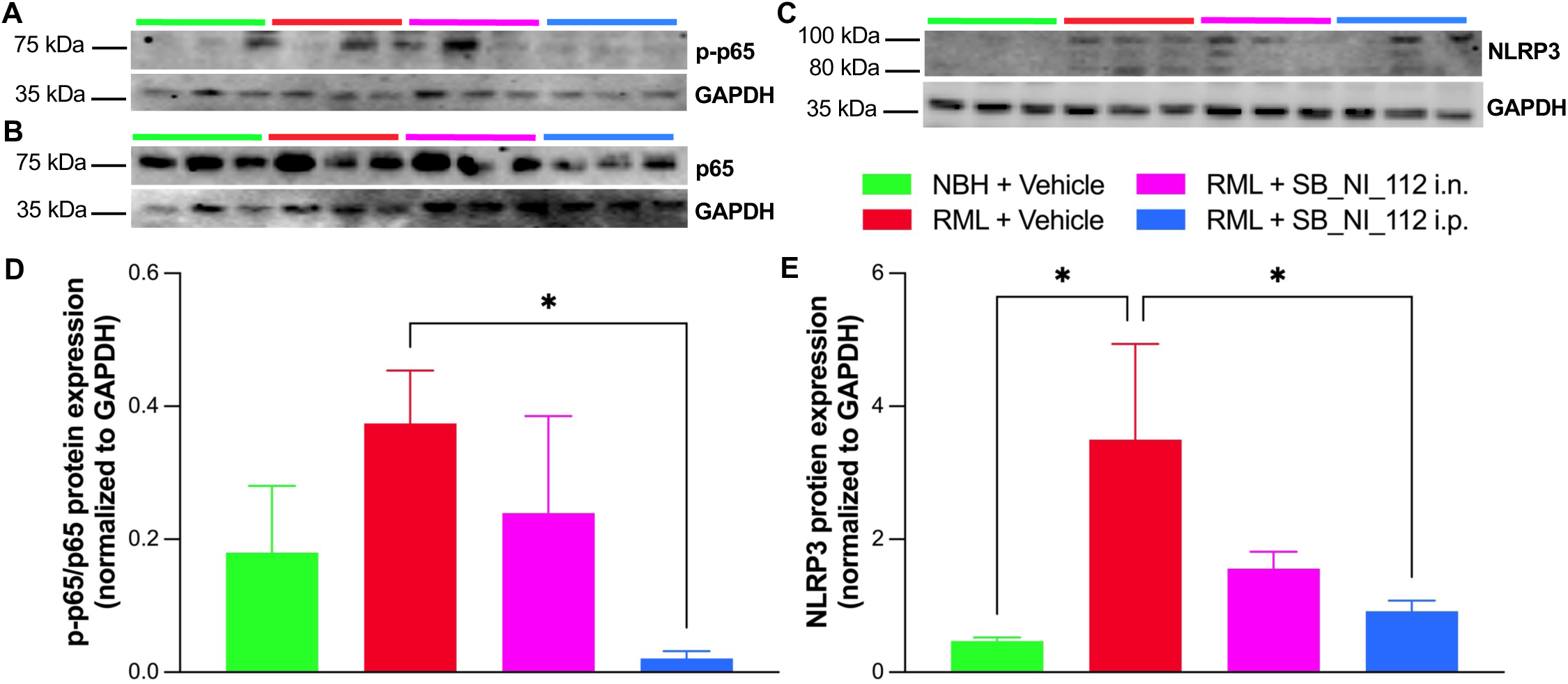
Intraperitoneally delivered SB_NI_112 significantly reduces phosphorylated p65 and NLRP3 protein expression in prion-diseased mice. Western blot images for phosphorylated-p65 (p-p65;A), p65 (B), and NLRP3 (C). Quantification of p-p65/p65 protein expression normalized to GAPDH (D). Quantification of NLRP3 protein expression normalized to GAPDH (E). N=3 for all groups. One-way ANOVA, error bars = SEM, * p< 0.05.

It is important to note that though many treatments for neurodegenerative diseases have been promising in early research, many fail due to toxicity and low efficacy rates in animal models or human trials. At this time, our research has identified no measurable toxicity or side effects associated with Nanoligomer SB_NI_112 at single high doses of 500 mg/kg or repeated dosing of 150 mg/kg. Additionally, the NF-κB and NLRP3 down-regulator is stable and bioavailable, has confirmed ability to cross the BBB, and displays high target specificity with minimal off-targeting. Our mechanism of action results (Figure 3) displaying superior inhibition of NF-|B and NLRP3 via i.p. route of delivery compared to i.n., combined with behavioral and histological data, support that an intraperitoneal dosing method is more successful at delivery therapeutic SB_NI_112 to the brain, thus providing better protection against neuroinflammation and prion disease progression.

A secondary study was completed at 150 mg/kg three times per week to explore the impact of a higher dose. However, we saw no difference in RML prion associated behavioral assessment scores and clinical signs of 10 mg/kg 2x per week and high dose 150 mg/kg 3x per week treatment groups (Figure 3 and Supplemental Figure 1 respectively), indicating there may be a threshold at which the treatment against neuroinflammation, specifically NF-κB and NLRP3, is effective against RML prion disease. Because there is no indication to treat with a higher dose, as it does not increase the therapeutic effect, we did not continue with additional methods to assess disease progression in the high dose study.

### Prion-induced behavioral and cognitive deficits occur significantly later in mice treated with NF-κB and NLRP3 down-regulator

Down-regulation of NF-κB and NLRP3 significantly protects against behavioral and cognitive changes associated with RML treated prion disease. RML mouse-adapted prion strain preferentially seeds and accumulates in the hippocampus^48,49^. Two common hippocampal behaviors affected by RML prion disease include nesting and burrowing activity^51,52^. To assess nesting, mice were given napkins to nest in overnight and scored between one (no nest formed) and five (fluffy built nest) weekly starting at 16 wpi. Monitoring of nest formation was found to be significantly higher among i.p. treated mice at 20 wpi (average score 4.5, p = 0.0001), 21 wpi (average score 3.2, p = 0.0007), and 22 wpi (average score, 2.5, p = 0.0216) (Figure 4A) compared to vehicle-treated prion-infected (average score 1 at 20 wpi, and 0.5 at 21 and 22 wpi) nest formation, while i.n. treated mice show a trend though no significant change in nesting throughout the disease. Similarly, a significant increase in the ability of the mice to build proper nests at 19 wpi (average score 4.5, p = 0.0039) and 21 wpi (average score 3.2, p = 0.0005) was shown with high dose i.p. treatment with SB_NI_112 compared to vehicle treatments (average score 2.5, p = 0.0039 at 19 wpi and 2.4, p = 0.0004 at 21 wpi) (Supplemental Figure 1A).

**Figure 4.**
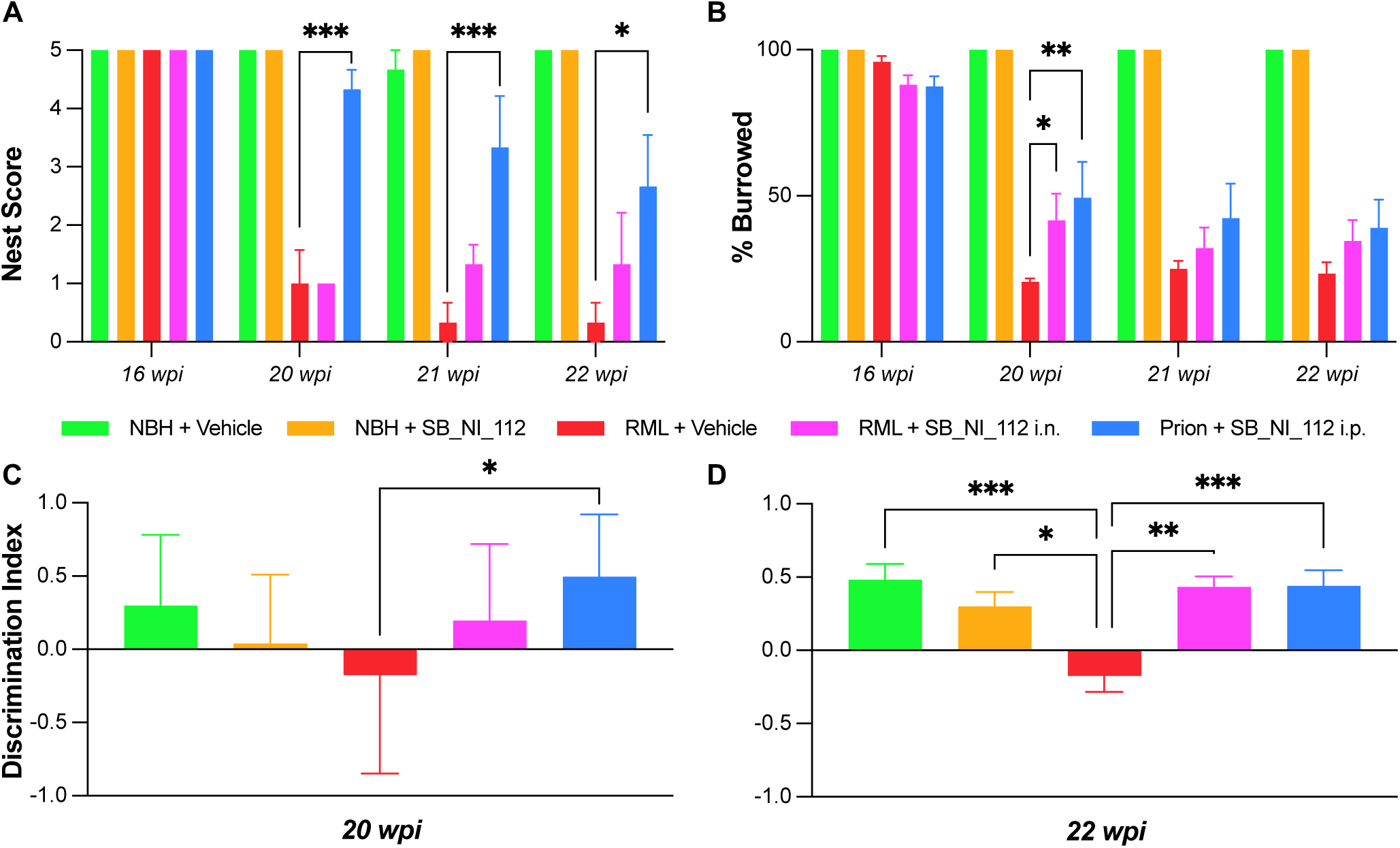
Behavioral and cognitive deficits are protected by NF-|B and NLRP3 down-regulation with SB_NI_112 in prion diseased mice. A significant increase in the ability of the mice to build proper nests at 20, 21, and 22 wpi with SB_NI_112 i.p. compared to vehicle treatments. Two-way ANOVA, error bars = SEM, * p < 0.05, ** p < 0.01, *** p < 0.001, **** p < 0.0001. In the hippocampal burrowing assay, i.p. and i.n. SB_NI_112 treated mice showed significantly better ability to burrow, along with high dose SB_NI_112 i.p. at 16 weeks, compared to untreated counterparts (B). Two-way ANOVA, error bars = SEM, * p< 0.05, ** p< 0.01. Only i.p. SB_NI_112 showed a significant difference in discrimination index at 20 wpi, higher discrimination index trends with i.n. SB_NI_112 (C). Both i.p. and i.n. SB_NI_112 treated mice had significantly higher scores in spatial novel object recognition at 22 wpi (D). N=9-12. One-way ANOVA with pos-hoc analysis, error bars = SEM, * p< 0.05, ** p< 0.01, *** p< 0.001, **** p< 0.0001.

Mice were given a plastic tube filled with food pellets within their home cages for 30 min to burrow freely. Following this time, the food pellets left in the tubes were weighed and the percentage burrowed was calculated. Prion-diseased mice are known to stop burrowing as the disease progresses^28,51,52^ but at 20 wpi, 21 wpi, and 22wpi, prion-diseased mice given therapeutic SB_NI_112 (i.n. and i.p.) had a later onset of deficits in burrowing activity (Figure 4B), showing a significance of protection at 20 wpi (average percent burrowed 47, p = 0.0475 for i.n.; average percent burrowed 42, p = 0.0032 for i.p.), trending through 22 wpi. (Figure 4B) A significant difference in the hippocampal burrowing assay was only indicated at 16 wpi (average percent burrowed was 75%, p > 0.0365) when mice were treated with the high dose SB_NI_112 compared to the vehicle (average percent burrowed was 51%) (Supplemental Figure 1B).

Cognitively, mice with RML-induced prion disease have a significant decrease in the ability to recognize a familiar object as the RML prion strain begins to aggregate within the hippocampus, the brain region essential for memory and learning^48,51,52^. Both i.n. and i.p. SB_NI_112 treatment protected prion-induced deficit in novel object recognition when compared to vehicle-treated diseased mice (Figure 4C, D). The ratio of time the mouse spent with the novel object to the familiar object (seen 24 hours beforehand) was calculated and noted by the discrimination index. The closer the discrimination index (DI) is to a value of one the more intact the hippocampal memory is, implying lower disease progression in the animal^53^. At 20 wpi, only SB_NI_112 i.p. showed a significantly higher discrimination index compared to untreated counterparts (mean discrimination index score of 0.4975, p = 0.0265), with a trend of improvement with low dose i.n. SB_NI_112 (mean discrimination index score of 0.1974, p =0.4383) (Figure 4C). At 22 wpi, both i.p. (mean discrimination index score of 0.4491, p = 0.0008) and i.n. treated mice (mean discrimination index score of 0.4343, p = 0.0022) had a significant increase when compared to vehicle-treated diseased mice with a discrimination index of -0.174 (i.e., preferring the familiar object) (Figure 4D). There was also a trend towards improved novel object recognition in high dose treated mice compared to untreated counterparts at 22 wpi (Supplemental Figure 1D).

### Down-regulation of NF-κB and NLRP3 with SB_NI_112 treatment reduces glial inflammation

Brain tissues were collected at 20 wpi and 24 wpi from low dose treatment groups. Formalin-fixed tissues were processed and embedded into paraffin wax and sectioned at 5µm. Brains were stained for both microglial and astrocytic inflammation (Figure 5-6). Microglia inflammation was identified as Iba1+ cells (Figure 5), and S100β+ cells denote inflamed astrocytes (Figure 6). Four main brain regions known to be affected in RML prion infection were analyzed: the hippocampus, cortex, thalamus, and cerebellum. Representative images of each brain region for all treatment groups are displayed for both microglial (Figure 5A-L) and astrocytic gliosis (Figure 6A-L), and arrows point out examples of positively stained cells.

**Figure 5.**
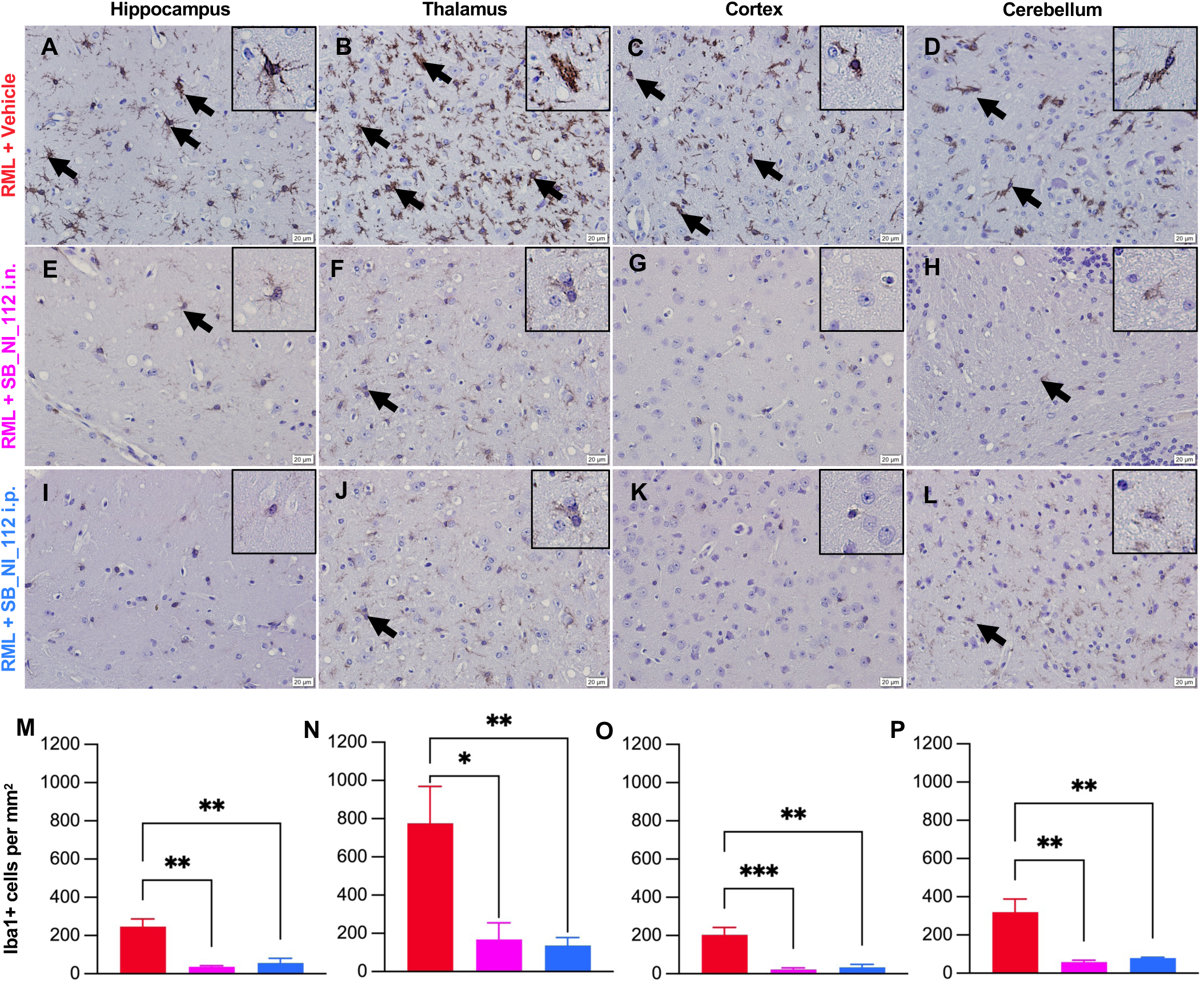
Microglial inflammation is significantly decreased by NF-κB and NLRP3 down-regulation with SB_NI_112 in prion diseased mice at 20 wpi. Representative images of the hippocampus, thalamus, cortex, and cerebellum with Iba1+ cells, a marker of activated microglia, with vehicle (A-D), SB_NI_112 i.n. (E-H), and SB_NI_112 i.p. (I-L) treatment. Quantitative analysis of each brain region identifies a significantly lower number of Iba1+ cells in the hippocampus (M), thalamus (N) cortex (O), and cerebellum (P) with both i.n. and i.p. treatment. Arrows represent examples of positive cells. Box insets show zoomed-in image of single microglia. Scale bar = 20µm, N=3-4. One-way ANOVA, error bars = SEM, *p > 0.05, **p> 0.001, ***p > 0.0001.

**Figure 6.**
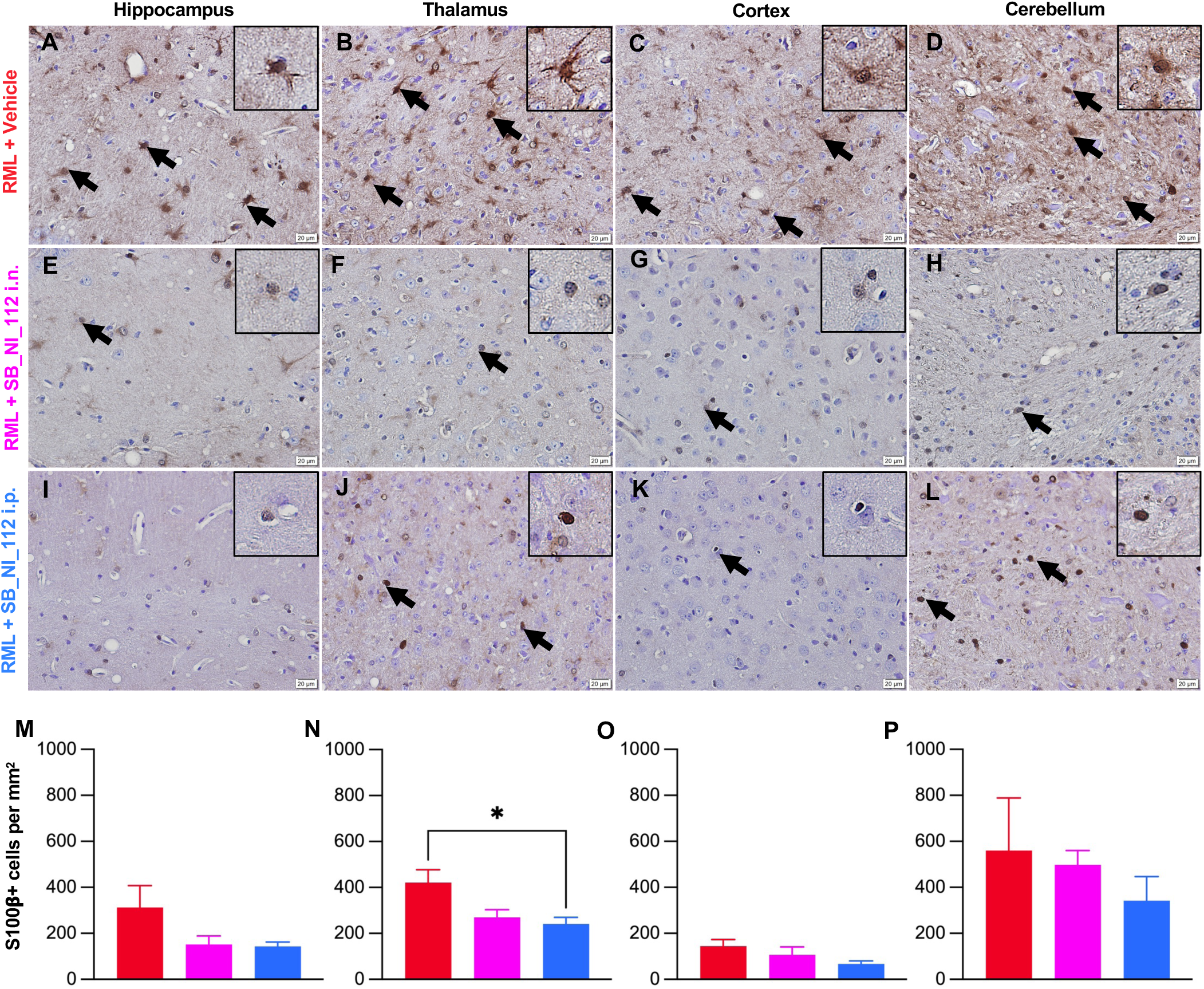
Astrocytic inflammation is significantly decreased by NF-κB and NLRP3 down-regulation with SB_NI_112 in prion disease mice at 20wpi. Representative images of the hippocampus, thalamus, cortex, and cerebellum with S100®+ cells, a marker of activated microglia, with vehicle (A-D), SB_NI_112 i.n. (E-H), and SB_NI_112 i.p. (I-L) treatment. Quantitative analysis of each brain region identifies a significantly lower number of S100®+ cells in the thalamus (N) and a trend of a decrease of inflamed astrocytes in the hippocampus (M) and cortex (O), and cerebellum (P) with treatment. Arrows represent examples of positive cells. Box insets show zoomed-in image of single astrocytes. Scale bar = 20µm, N=3-4. One-way ANOVA, error bars = SEM, * p< 0.05.

In the thalamic region, both microglial and astrocytic inflammation were significantly suppressed by down-regulation of NF-κB and NLRP3 with SB_NI_112, both i.n. and i.p. (Figure 5N and 6N respectively) at 20 wpi. Within the thalamic region, vehicle treated samples had a mean of 775.4 Iba1+ microglia per mm^2^, followed by SB_NI_112 i.p. treatment with a mean of 168.6 Iba1+ microglia per mm^2^ (p = 0.0115 compared to vehicle), and SB_NI_112 i.n. treatment with a mean of 168.6 Iba1+ microglia per mm^2^ (p = 0.0087 compared to vehicle). In regard to astrogliosis in the thalamus, vehicle treated samples had a mean of 421 S100β+ astrocytes per mm^2^, followed by SB_NI_112 i.p. treatment with a mean of 270 S100β+ astrocytes per mm^2^, (p = 0.0613 compared to vehicle), and SB_NI_112 i.n. treatment with a mean of 240.8 S100®+ astrocytes per mm^2^, (p = 0.0282 compared to vehicle). Within the hippocampal region, vehicle treated samples had a mean of 246 Iba1+ microglia per mm^2^, followed by SB_NI_112 i.n. treatment with a mean of 56.31 ba1+ microglia per mm^2^ (p = 0.0019 compared to vehicle), and SB_NI_112 i.p. treatment with a mean of 36.75 Iba1+ microglia per mm^2^ (p = 0.0010 compared to vehicle). In the cortex, vehicle-treated samples had a mean of 203 Iba1+ microglia per mm^2^, followed by SB_NI_112 i.n. treatment with a mean of 33.22 Iba1+ microglia per mm^2^ (p = 0.0014 compared to vehicle), and SB_NI_112 i.p. treatment with a mean of 22.32 Iba1+ microglia per mm^2^ (p = 0.0010 compared to vehicle). Lastly, in the cerebellum vehicle-treated samples had a mean of 319.6 Iba1+ microglia per mm^2^, followed by SB_NI_112 i.n. treatment with a mean of 80.05 Iba1+ microglia per mm^2^ (p = 0.0023 compared to vehicle), and SB_NI_112 i.p. treatment with a mean of 57.71 Iba1+ microglia per mm^2^ (p = 0.0013 compared to vehicle). There was a trend towards astrocyte reduction for both i.p. and i.n. treatment groups in the hippocampus (Figure 6M), Cortex (Figure 6O), and cerebellum (Figure 5P).

Microglia are closely associated with plaques in multiple forms of human prion disease including Creutzfeldt-Jakob disease, Gerstmann-Straussler-Scheinker, and kuru and it has been shown that microglia play a role in neuronal cell death during prion disease^54,55^. Microglial activation appears before the manifestation of clinical signs of prion disease, indicating microglial inflammation plays a role in the progression of prion disease. Furthermore, the deposition of prion proteins alone does not directly predict regions of neurodegeneration^55^. In addition to microglial activation, astrogliosis is found early in prion disease pathology, and reactive or inflamed astrocytes are known to be synaptotoxic in the pathogenesis of prion disease^22,23,56,57^. Further, NF-κB and NLRP3 activation are implicated in the induction of neurotoxic astrocytes by microglia. Global microglial ablation of NLRP3 ameliorates A1-like astrocytes in vitro and in vivo, decreasing neuronal dysfunction. Our research supports this claim by displaying decreased microglial and astrocytic inflammation down-regulating NLRP3 and NF-|B with brain-penetrant SB_NI_112 (Figure 5-6). Though a less significant difference was seen in inflamed astrocytes (S100β+ cells) between treated and untreated groups, this may be due to temporal differences in microglial and astrocyte activation throughout chronic inflammation or a proposed cell-type specific activation of NLRP3 in microglia compared to astrocytes^58,59^.

### SB_NI_112 protects against prion-induced spongiosis and neuronal loss

Morphologic spongiosis or vacuoles within the brain tissue is a hallmark of prion disease that increases as the disease progresses^44,45^. To assess and quantify this change, we used pathological scoring of the hippocampus, thalamus, cortex, and cerebellum following H&E staining. Scores were given ranging from + (mild spongiosis) to +++ (high spongiosis) by three blinded researchers Figure 6E-P. This revealed protection against spongiform change in both i.p. and i.n. SB_NI_112 treatment of prion diseased mice when compared to vehicle-treated prion diseased mice at 20 wpi. Representative images from each brain region with corresponding average spongiosis score are displayed for RML + Vehicle, RML + SB_NI_112 i.n., and RML + SB_NI_112 i.p. Overall, down regulation of NF-κB and NLRP3 with SB_NI_112 protects against hallmark spongiosis in prion disease. Beyond spongiosis, the loss of neurons within the hippocampus following RML prion infection is well established^48,49^. Therefore, we quantified neuronal loss in SB_NI_112 treatment groups to untreated counterparts. Neuronal numbers within the CA1 region of the hippocampus (Figure 7A-C) were significantly higher with down-regulation of NF-|B and NLRP3 by low dose i.n. and i.p. delivered SB_NI_112 compared to the vehicle (Figure 7D, p>0.05). Critically, SB_NI_112 prevents the neuronal loss and spongiform change induced by RML prion infection. As neuronal loss is the direct cause of behavioral and cognitive deficits, and ultimate fatality due to prion disease, it is imperative potential therapeutics show success in this metric^49^.

**Figure 7.**
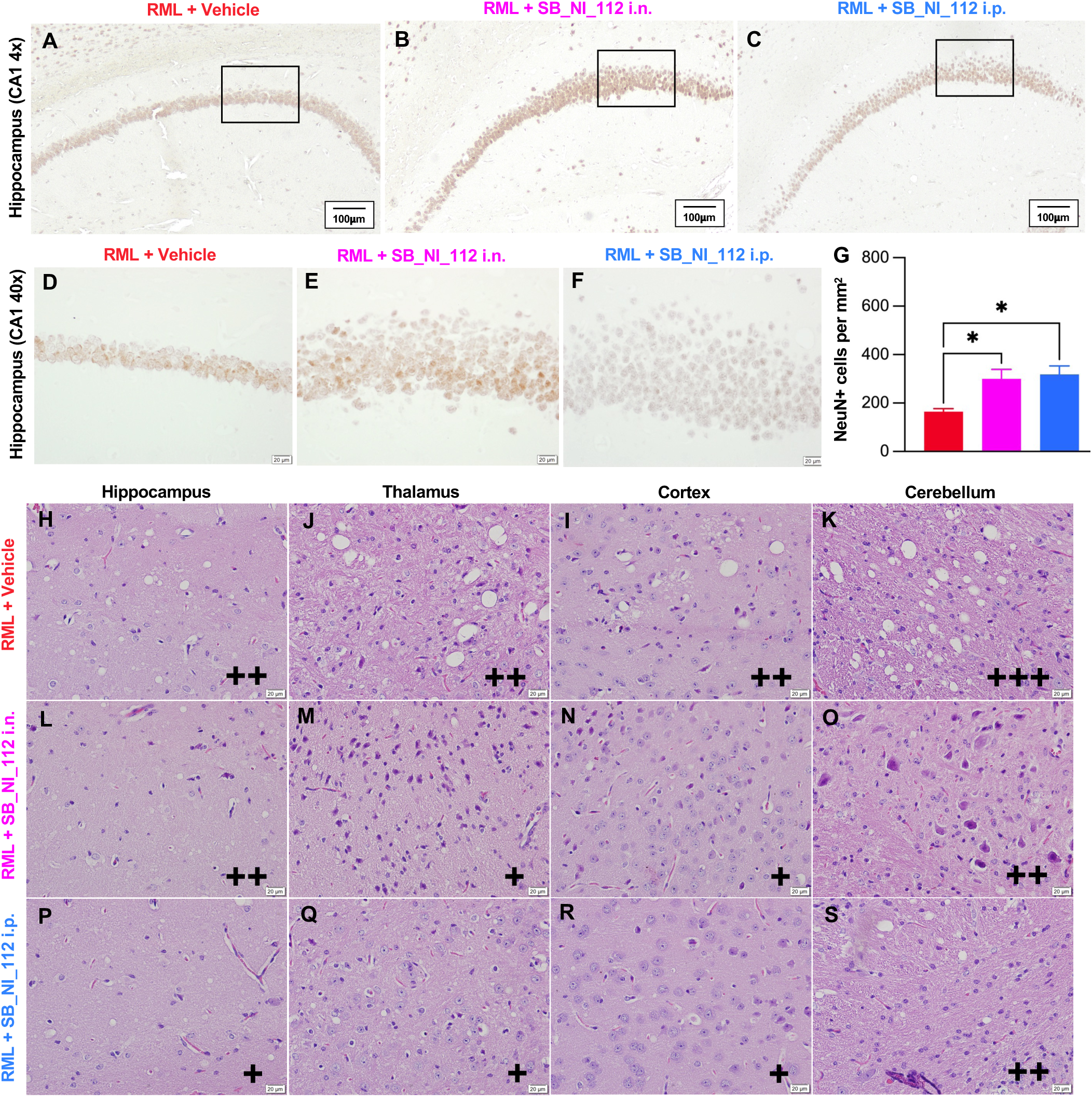
Prion induced neuronal loss and spongiotic change is significantly decreased by NF-κB and NLRP3 down-regulation with SB_NI_112 in prion diseased mice. Representative images of the NeuN + cells (neuronal cell bodies) in the CA1 region of the hippocampus (A-C) Significant decrease in the number of neurons lost at 20 wpi with treatment via i.p. and i.n. SB_NI_112 (D). Scale bar = 20mm, N=3-4. One-way ANOVA *p > 0.05. Representative images of the spongiotic change in hippocampus, thalamus, cortex, and cerebellum with vehicle (E-H)), low dose SB_NI_112 i.n. (I-L), and low dose SB_NI_112 i.p. (M-P). Pathological scoring of each brain region identifies a protection or decrease of pathological score at all brain regions analyzed; + = mild spongiosis, ++ = moderate spongiosis, +++ = severe spongiosis. N=3-4.

### Treatment with SB_NI_112 improved clinical scores and survival in mice despite accumulation of PrP^Sc^

At 22 wpi, vehicle-treated mice begin to show a significant increase in clinical signs of prion disease (Figure 8A). From 22 wpi to 24 wpi i.p. SB_NI_112 treated diseased mice have significantly (p<0.0001 for all weeks) lower clinical scores than vehicle-treated diseased mice, and SB_NI_112 i.n. treatments displayed significantly lower scores at 22 wpi (p <0.0001) and 23 wpi (p = 0.0197). Untreated Prion clinical signs included tail rigidity, hyperactivity, ataxia, extensor reflex (clasping), tremors, righting reflex, kyphosis, and poor grooming^28,48,52^. Each sign was rated on a scale from 0-5. All clinical sign scores were combined for a total score. SB_NI_112 i.p. treated mice had clinical scores with a mean lower than 2 at 22 wpi, 3 at 23 wpi, and 5 at 24 wpi, compared to clinical score mean of 6+ points higher in untreated counterparts (Fig 8A). Mice receiving i.n. treatments averaged scores of 3 at 22 wpi and 7 at 23 wpi (Fig 8A). Of note, clinical scores in high dose treated mice were significantly lower (p = 0.0120) at 22 wpi, with a trend towards lower scores at 23 and 24 wpi (Supplemental Figures 1E). See supporting videos for visual of clasping (supporting videos 1-3), righting reflex (supporting videos 4-6), and walking (supporting videos 7-9) in Control (NBH), RML + vehicle, and RML + Sb_NI_112 i.p. at 22 wpi. Mice were considered terminal and euthanized after reaching a total score of 9 or above. Additionally, mice with a weight loss greater than 10% were considered terminal and euthanized. The date of euthanasia was documented for survival analysis (Figure 8B).

**Figure 8.**
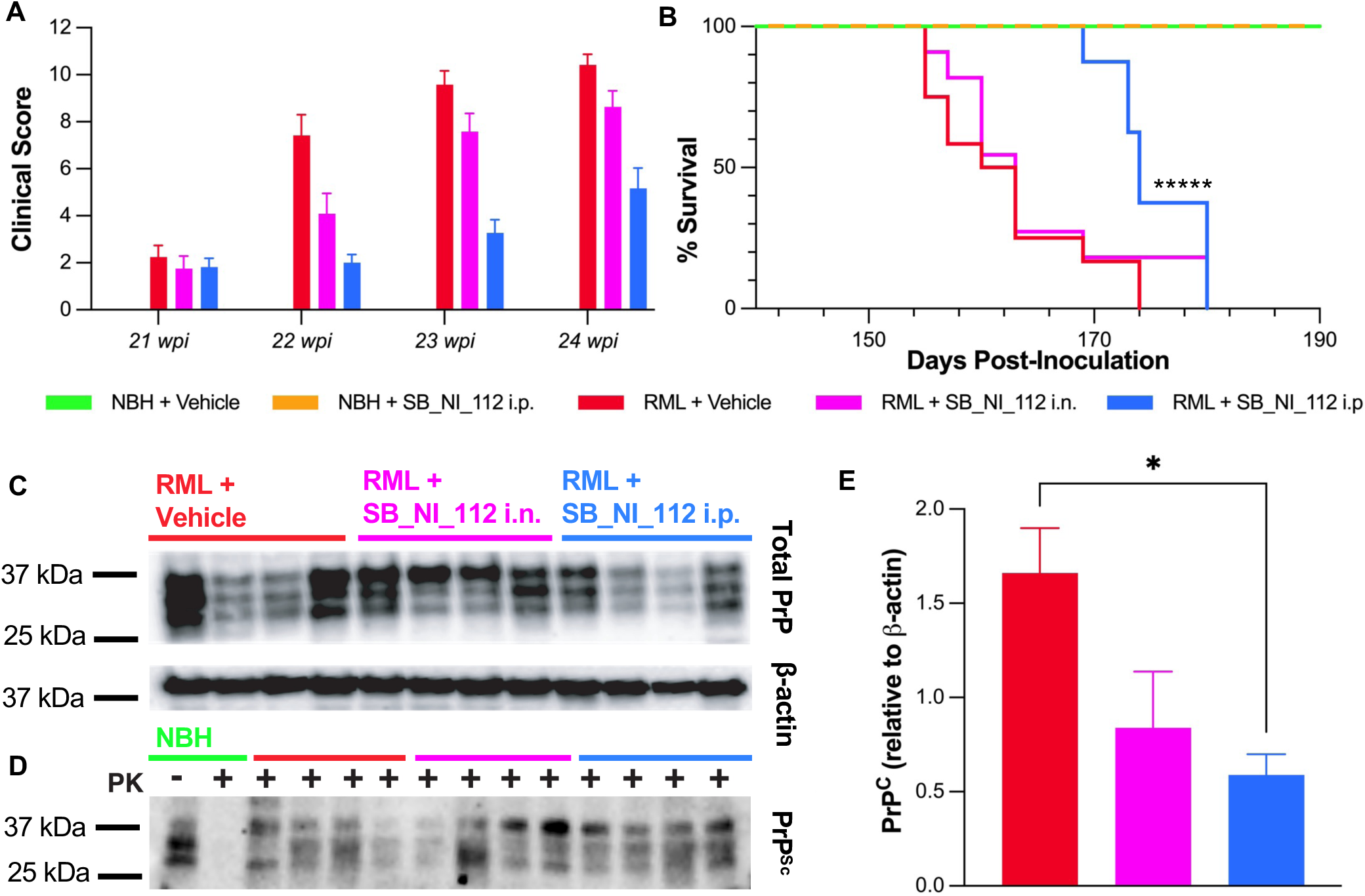
Prion diseased mice treated with NF-κB and NLRP3 down-regulator had significantly lower clinical scores and increased lifespan, independent of PrP^Sc^ accumulation. Clinical scores of mice treated with i.n. or i.p. SB_NI_112 at 22 and 23 wpi were significantly lower than vehicle treated counterparts, continued significance at 24 wpi with i.p. SB_NI_112 (A). N=9 for prion positive i.p. and i.n. groups, N=9-12 for all other groups. One-way ANOVA, error bars = SEM, * p < 0.05, ** p < 0.01, *** p < 0.001, **** p < 0.0001. Mice treated with low dose SB_NI_112 i.p. lived significantly longer than untreated counterparts (B) Survival curve, ***** (p <0.000001). Total PrP and PrP^Sc^ levels detected after proteinase K (PK) (indicates not added, + indicates added) digestion at 20 wpi measured by immunoblot (C). Total PrP relative to β-actin was significantly different among SB_NI_112 i.p. treated and untreated groups (D). N=4 for all groups. One-way ANOVA, error bars = SEM, * p < 0.05. No significant difference was seen in total PrP^Sc^ relative to PrP^C^ (E). Total PrP^Sc^ relative to PrP^C^ was equivalent in prion-infected mice treated with vehicle and SB_NI_112 i.n. and i.p. treatment (E). N=4 for all groups. One-way ANOVA, error bars = SEM. All immunoblots were performed on frontal cortex lysates.

The lifespan of prion disease mice was significantly increased with i.p. SB_NI_112 Nanoligomer treatment (p <0.000001) compared to vehicle-treated mice (Figure 8B). No change or protection of i.n. delivered SB_NI_112 was seen in survival of prion-diseased mice. Human sporadic Creutzfeldt-Jakob disease has a mean survival of 7-8 months after diagnosis^60^. On average, i.p treatment with SB_NI_112 significantly increased prion-diseased mouse lifespan. Untreated mice had a mean survival of 162 days, i.p. SB_NI_112 treated 174 days, and i.n. SB_NI_112 163 days. Of note, the first mouse without SB_NI_112 treatment reached a clinical score indicative of sacrifice 15 days before the first mouse receiving i.p. treatment. 15 mouse days equate to approximately 6 months in human years, a near doubling of expected survival time after diagnosis^61^. Importantly, there was no toxic pathology induced by the down regulation of NF-|B and NLRP3 by SB_NI_112 (Figure 2A).

To determine if SB_NI_112 modulates the accumulation of the misfolded prion protein, PrP^Sc^, immunoblotting was performed on cortical brain homogenates for levels of the cellular PrP^C^ and proteinase K-resistant PrP^Sc^ (Fig 8C, Figure S4). Total PrP relative to β-actin was significantly different among treated and untreated groups (Figure 8D). No significant difference was seen in total PrP^Sc^ relative to PrP^C^ (Figure 8E, Figure S4). Similarly, other studies examining potential prion therapeutics have shown similar results in regard to total PrP^Sc48,49^. This indicates dampening chronic inflammatory signaling protects against prion-associated neurodegeneration independent of PrP^Sc^ accumulation. SB_NI_112, down-regulator of NF-κB and NLRP3, shows efficacy in slowing the disease clinically, behaviorally, and pathologically while increasing predicted survival after diagnosis, and use as a combination therapy for individuals diagnosed with prion disease.

### NLRP3 inflammasome involvement in progression of prion disease

The level of influence the NLRP3 inflammasome has on prion disease progression is still debated within the literature. The NLRP3 inflammasome was found to be involved in lipopolysaccharide (LPS)-primed microglia after exposure to a synthetic neurotoxic prion fragment in vitro^29,30^. Previous studies highlight the importance of the NLRP3 inflammasome in the production of IL-1β and IL-18 in systemic inflammation and neurodegenerative disease, including prion disease^21,24–26,62^. An *in vitro* study in BV2 microglia and co-cultured glia reported that aggregated prion protein was involved in both priming and activation of the NLRP3 inflammasome^29^.

Regarding cytokine production in our study, at 16wpi, IL-28 is significantly reduced in i.p. treated (average 30.1 pg/mg total protein, p <0.05) mice compared to untreated counterparts (average 43.4 pg/mg total protein) (Figure S2). Interestingly, an *in vivo* study found levels of IL-1β at end-stage prion disease were not affected by the absence of NLRP3 and that NLRP3 knockout mice succumbed to RML prion in a similar time frame to wild-type counterparts^63^. Conversely, our *in vivo* research supports the involvement of the NLRP3 inflammation in prion disease progression, as downregulation of NLRP3 and upstream signaling molecule NF-|B decreased glial inflammation, protected against neuronal loss, prevented spongiotic change, rescued cognitive deficits, and significantly lengthened lifespan of prion-diseased mice. Additionally, inflammatory cytokines IL-2 (Figure S2 B), IL-1β (Figure S2 C), and IL-1α (Figure S2 D), trended lower in prion-diseased mice treated with i.p. SB_NI_112, like NBH controls, when compared to untreated counterparts. Principal component analysis revealed trending differences between treated and untreated prion-diseased mice (p = 0.094) (Figure S2 E). Based on conflicting data, there are likely additional inflammatory pathways upregulated by PrP^Sc^ in conjunction with the known direct degenerative impact PrP^Sc^ aggregates have on neurons^64^.

Limitations of our study include a low sample size at each time point. Higher significance may have been achieved with a larger sample size, particularly in regard to later behavioral testing timepoints. Moreover, adding more behavioral time points (weekly vs. biweekly) could display more detailed changes in behavior as prion disease progresses. Additionally, there may be other routes of dosing that provide stronger protection against prion disease progression, including intravenous, subcutaneous, or oral routes. Finally, because the therapeutic showed high safety efficacy, it would be beneficial to test a variety of difference dose concentrations and frequencies.

### Conclusions and Future Prospects

Down-regulation of NF-κB and NLRP3 in a mouse-adapted RML prion model slows disease progression by reducing glial inflammation, delaying development of behavioral deficits and clinical signs, preventing loss of neurons, and extending lifespan. Because treatments successful in prion disease are anticipated to work in other neurodegenerative diseases due to common pathogenesis, these findings, if validated clinically, have important implications for patients diagnosed with prion disease and those suffering from more widespread neurodegenerative diseases like AD and PD.

Though all mice eventually succumbed to disease in this study, PrP^Sc^ misfolding and accumulation in prion disease, in comparison to other misfolded proteins in neurodegenerative diseases (Aβ in AD and α-Syn in PD), is substantially more rapid and aggressive^65^. Additionally, properly functioning microglia can phagocytose and degrade both Aβ and α-Syn. It is only when microglia lose the ability to degrade misfolded proteins and the load of Aβ or α-Syn becomes too high that aggregation and progression of disease state occurs in AD and PD respectively^24–27^. Recent research supports chronic inflammation and NLRP3 inflammasome stimulation decreases microglial autophagy, leading to increased Aβ or α-Syn production and aggregation in the brain^39–41^. Furthermore, inflammation in other glial cells, namely astrocytes, leaves them unable to properly perform metabolic, structural, homeostatic, and neuroprotective tasks^65^. Our study revealed down-regulation of NF-κB and NLRP3 decreases astrocyte inflammation in addition to microglial inflammation.

Specifically in prion disease, there have been few small molecules, antibodies and other compounds that show promise in treatment of disease pathogenesis. Therapeutic oligonucleotides for the prion protein or PrP-lowering antisense oligonucleotides (ASO) have proven to lower the toxic load of the disease-causing misfolded prion protein increasing survival of infected mice. Critically this was irrespective of the strain or type of prion the mice were exposed to but, only successful with the highly invasive intracerebroventricular injection, a critical limitation of this treatment^66^. The use of antibodies as an immunotherapy and small molecules against the prion protein has been studied for years and in rodents the progression and acceleration of the toxic PrP^Sc^ is reduced, but unfortunately the protection of these do not translate well to patients^47,67–72^. The development of human specific antibodies is showing some efficacy in patients however the need for the ability to give the immunotherapy to patients earlier, prior to severe disease states, seems to be essential^73^.

Nanoligomers present benefits compared to other small-molecules, antibodies, and antisense oligonucleotide (ASO) platforms. First, their efficient biodistribution and facile delivery across the BBB and reaching various sections of the brain without using invasive administration methods such as Intracerebroventricular injection used in other ASO based treatments.^66,74^ Second, lack of any immunogenic response, any adverse systemic effect or accumulation makes these molecules preferable to other small molecules. This study focusing on a Prion infection model, showed that Nanoligomers directed at the inflammasome demonstrate neuroprotection, improved animal survival rates, and enhanced cognitive function. This underscores the technology’s safety and broader potential applications. Another significant advantage of the Nanoligomers discovery engine lies in its capability to selectively target specific gene(s) for either upregulation or downregulation of expression as needed. Notably, recent research demonstrated that a low dosage of 5 mg/kg of Nanoligomer significantly reduced inflammation in the mouse hippocampus during LPS-induced neuroinflammation.^42^

Limitations of our study include a low sample size at each individual time point. Higher significance may have been achieved with a larger sample size, particularly regarding behavioral testing timepoints for the later more diseased timepoints. Moreover, adding more behavioral time points (weekly vs. biweekly) could display more detailed changes in behavior as prion disease progresses. Additionally, there may be other routes of dosing that provide stronger protection against prion disease progression, including intravenous, subcutaneous, or oral routes. Finally, because the therapeutic showed high safety profile, it would be beneficial to test a higher dose concentration and more often frequency in dosing. In summary, targeting the NLRP3 inflammasome via down-regulation of NF-|B and NLRP3 with SB_NI_112 has promising therapeutic potential for neurodegenerative diseases. SB_NI_112 reduces inflammatory signaling amongst microglia and astrocytes, which is highly neuroprotective. At the least, targeting chronic neuroinflammation in neurodegenerative diseases is an essential component in the “cocktail” of multi-target therapeutic interventions that are likely necessary for successful modification and halting of disease.

## Methods

### Mice and Brain homogenates

C57Bl/6 (Jackson Laboratory) mice were intracranially inoculated with 30µl of 0.1% 22L or Rocky Mountain Laboratories (RML) strains of mouse-adapted prions, or normal brain homogenate (NBH), in 1% Pen-Strep (1:1 penicillin-streptomycin). Mice were monitored for weight loss and clinical signs of prion disease and euthanized after showing signs of terminal illness. 20% brain homogenates in phosphate-buffered saline (PBS) were made using beads and a tissue homogenizer (Benchmark Bead Blaster 24) and stored at -80C. Brain homogenates were aliquoted and treated with UV light for 30 minutes to sterilize before being used for cell culture.

### Nanoligomer Design and Synthesis

The Nanoligomer design, sequences, and synthesis has been described in our previous work (59). Nanoligomers were designed and synthesized (Sachi Bioworks Inc.) according to published methods^42,43,50,75,76^. The Nanoligomer is composed of a nanobiohybrid molecule based on antisense peptide nucleic acid (PNA)^42,43,75,77,78^ conjugated to nanoparticle^1,2,5^ for improved delivery. The PNA moiety was chosen to be 17 bases long (to optimize solubility and specificity) antisense to the start codon regions of nfkb1 (N terminus- -C terminus, sequence: AGTGGTACCGTCTGCTA) and nlrp3 (N terminus- -C terminus, sequence: CTTCTACTGCTCACAGG) within the Mouse genome (GCF_000001635.27).^1^ Potential PNA sequences were screened for solubility, self-complementing sequences, and off-targeting within the mouse genome (GCF_000001635.27). The PNA portion of the Nanoligomer was synthesized on Vantage peptide synthesizer (AAPPTec, LLC) with solid-phase Fmoc chemistry. Fmoc-PNA monomers were obtained from PolyOrg Inc., with A, C, and G monomers protected with Bhoc groups. Following synthesis, the peptides were conjugated with gold nanoparticles and purified via size-exclusion filtration. Conjugation and concentration of the purified solution was monitored through UV-Vis (Thermo Scientific, NanoDrop) measurement of absorbance at 260 nm (for detection of PNA) and 400 nm (for quantification of nanoparticles). Nanoligomers were stored at 4 °C until further use.

### Biodistribution and Toxicity

Six female and six male C57Bl/6 (Jackson Laboratory) mice, 8-10 weeks of age, were injected with a single dose of 500 mg/kg intraperitoneal or intranasal SB_NI_112. Mice were monitored for 2 weeks for safety/tox study and dosing. Weight was monitored daily, along with any changes in behavior/well-being. At the end of the study, mice were euthanized, and tissues harvested for histopathological scoring, blood concentration of any inflammatory markers, albumin content in serum. Using three-fold lower concentration than the maximum tolerated dose (MTD) (150 mg/kg), eight female and eight male C57Bl/6 mice were dosed for 2 weeks with 2 doses/week. Mice were euthanized and harvested to measure distribution of Nanoligomer through different tissues using ICP-MS.

### Nanoligomer Administration

At 10 weeks post RML prion or NBH inoculation, mice were treated two times per week either intraperitoneally (i.p.) or intranasally (i.n.) with 10 mg/kg of SB_NI_112 diluted in sterile phosphate buffered saline (PBS). For high dose study, mice were treated three times per week with 150 mg/kg of SB_NI_112 i.p. Mice receiving treatment i.n. were anesthetized prior to administration and laid in an intranasal apparatus that controlled isoflurane throughout the intranasal procedure.

### Cognitive assay: Novel Object Recognition

Mice were tested in a rectangular arena (28cm x 43cm) and habituated by exploring the empty arena for ten minutes. During day one of testing, mice were placed in the arena with two identical objects (constructed with Legos) and video monitored for 5 minutes of exploration. On day two of testing, one of the known objects was replaced by a novel object, and the 5-minute exploration was filmed using a 1080P FHD Mini Video Camera. All objects and the arena were cleansed thoroughly with 70% ethanol between trials to ensure the absence of olfactory cues. Time exploring both the known and novel object was calculated by blinded observers. Discrimination index (DI) calculated following Absolute vs Relative Analysis protocol^53^. All objects were tested prior to use to ensure the mice would explore them.

### Hippocampal Behavioral assays: Burrowing and Nesting

Briefly, mice were placed in a large cage with a PVC tube (2.33” x 5.75”) full of food pellets, as describe^28,48,51^. The percentage of burrowing activity is calculated from the difference in the weight of pellets in the tube before and after 2 hours. To test nesting, we placed three fresh napkins in each cage. After 24 hours, cages were examined for nesting activity. Nests were scored on a scale of 0-5, where 0 represents no nesting activity, and 5 represents a high-quality nest.

### Clinical Scoring of mice

Eight key clinical signs were monitored in mice daily beginning at 20 or 21 wpi. Clinical signs included tail rigidity, hyperactivity, ataxia, extensor reflex, tremors, righting reflex, kyphosis, and poor grooming. Each sign was rated on a scale from 0-5. All clinical sign scores were combined for a total score. Mice considered terminal and euthanized after reaching a total score of 9 or above. Additionally, mice with a weight loss greater than 10% were considered terminal and euthanized.

### Immunohistochemistry

Paraffin-embedded brains were sectioned at 4 µm and stained with NeuN antibody (1:250; Cell Signaling) for CA1 neuronal counts. Astrogliosis was detected with S100β antibody (1:400; Abcam ab41548) and Iba1 antibody (1:400; Abcam, ab5076) was used for microglial counts. Nonspecific binding was blocked before primary antibodies with 10% Horse Serum (Sigma Aldrich). A biotinylated secondary antibody (Vector Labs) was used, and stain was visualized with diaminobenzidine reagent. All images were taken with CellSens Dimension Desktop 3.1 and counted with CellSens (Iba1 and S100β+) or manually (NeuN).

### Immunoblotting

Cell lysates were isolated using the protein lysis buffer (50mM Tris, 150mM NaCl, 2mM EDTA, 1mM MgCl2, 100mM NaF, 10% glycerol, 1% Triton X-100, 1% Na deoxycholate, 0.1% SDS and 125mM sucrose) supplemented with Phos-STOP and protease inhibitors (Roche). A BCA Protein Assay kit (Thermo Scientific) was used to quantify protein concentration of lysates, and 250 mg or 500mg protein was digested with 20 µg/ml proteinase K (PK) (Roche) for PrP^Sc^ blots for 1 hour at 37C. This digestion allows all other proteins to be digested so PrP^Sc^ can be detected, therefore loading control antibodies cannot be used for these blots. Digestion was terminated with 2mM PMSF and lysates were spun at 40,000 x g for 1 hour at 4C before being loaded on a gel. For PrP^C^ blots, 20 mg of samples was used. Samples were run using 4-20% acrylamide SDS page gels (BioRad) and then transferred onto 0.45µm PVDF blotting paper (MilliPore). Primary antibody Bar-224 (Cayman Chemical Company) was used at 1:1,000 dilution for PrP^Sc^ blots and 1:5,000 dilution for PrP^C^ blots. HRP-conjugated secondary antibodies were used at a concentration of 1:5,000 (Vector Laboratories). For PrP^C^ blots, loading control GAPDH was ran at a 1:5,000 dilution (MilliPore), with HRP-conjugated anti-mouse secondary at 1:5,000 dilution (Southern Biotech). The protein antibody complex was visualized using SuperSignal West Pico PLUS Chemiluminescent Substrate (Thermo Scientific) and visualized with the BioRad ChemiDoc MP.

### Quantification of Cytokine & Chemokine in mouse tissues

We used previously published protocols (58-59). Briefly, dissected brain tissues were flash-frozen and homogenized using a mortar and pestle to form a homogenate solution in tissue cell lysis buffer (Thermo Fisher, EPX-99999-000). Samples were then centrifuged at 16,000xg for 10 min at 4°C and supernatant was transferred to a fresh tube. Bio-Rad’s DC Protein Assay Kit was used to determine protein content and all samples were diluted to 10 mg protein/mL. Quantification of cytokines/ chemokine was performed with Cytokine & Chemokine 36-Plex Mouse ProcartaPlex Panel 1A (EPXR360-26092-901), analyzed on a Luminex MAGPIX xMAP instrument, and quantified using xPONENT software. For standard curves, eight four-fold dilutions of protein standards were used.

### ICP-MS

Briefly, tissue samples were homogenized in a TissueLyser bead mill (Qiagen, Germantown, MD) at 30 Hz for 3 min with the addition of 1 μl deionized water per 1 mg organ tissue. Volumes of homogenates ranging from 10 to 130 μl were then digested in 500 μl of aqua regia (3:1 hydrochloric acid to nitric acid) for 4 hours at 100 °C. Pellets were resuspended in water and analyzed with a NexION 2000B single quadropole ICP-MS (PerkinElmer, Waltham, MA). A Meinhard nebulizer was used with a cyclonic glass spray chamber for the introduction of the sample. A nickel sample and skimmer cone were used with an aluminum hyperskimmer cone. The ICP-MS was optimized daily with a calibration solution of 1 ppb In, Ce, Be, U, and Pb. Au was the analyte monitored. Data was collected using the sample acquisition module in Syngistix software (version 2.3). Data was analyzed using Microsoft Excel by converting measured Au concentrations to pg/mg organ via organ homogenate volume used. A seven-point linear curve with a 1000 ppm Au standard solution (Ricca Chemical Co, Arlington, TX) was generated at a concentration range of 0-250 ppb. Indium was spiked into each sample at a concentration of 5 ppb. Linearity of the standard curve was defined at an R^2^ value of greater than 0.995. TraceMetal grade 70% HNO3-was purchased from ThermoFisher Scientific (Waltham, MA). All H2O used was from a Milli-Q system (Millipore, Burlington, MA).

### Statistical Analysis

All data is presented as mean +/- SEM unless otherwise specified. A ROUT (Q = 10%) outlier test was performed on all datato identify any potential outliers, which were removed from the data set. Difference between experimental groups were analyzed using a One-way ANOVA or Two-way ANOVA. Statistical analysis was completed using Prism. Significance is denoted throughout the manuscript as *= p ≤ 0.05, **= p ≤ 0.01, ***= p ≤ 0.001 and ****= p ≤ 0.001.

### Ethics Approval

Mice were euthanized by deeply anaesthetizing with isoflurane followed by decapitation. All mice were bred and maintained at Lab Animal Resources, accredited by the Association for Assessment and Accreditation of Lab Animal Care International, in accordance with protocols approved by the Institutional Animal Care and Use Committee at Colorado State University.

## Acknowledgments

The authors thank Colorado State University Lab Animal Resources for their animal husbandry.

## Abbreviations

AD: Alzheimer’s Disease
PD: Parkinson’s Disease
NF-|B: Nuclear factor kappa B
NLRP3: NOD, LRR, and pyrin-containing 3
wpi: weeks post inoculation

## Author Contributions

Conceptualization: JAM, PN, AC, SM

Methodology: SJR, SWB, SS, GMW, PS, ADH, AJDH, VG

Funding acquisition: JAM, PN and AC

Supervision: JAM, PN, AC

Writing – original draft: SJR, JAM

Writing – review & editing: JAM, PN, AC, SM, SJR

## Conflict of Interest

The authors declare the following competing financial interest(s): SS, AC, and PN are employed by Sachi Bioworks where this technology was developed. AC and PN are the founders of Sachi Bioworks, and PN has filed a patent on this technology.

## Funding Sources

NASA SBIR 80NSSC22CA116, Sachi Bio, and CVMBS Murphy Turney Fund at Colorado State University

---

**Supporting Information.**
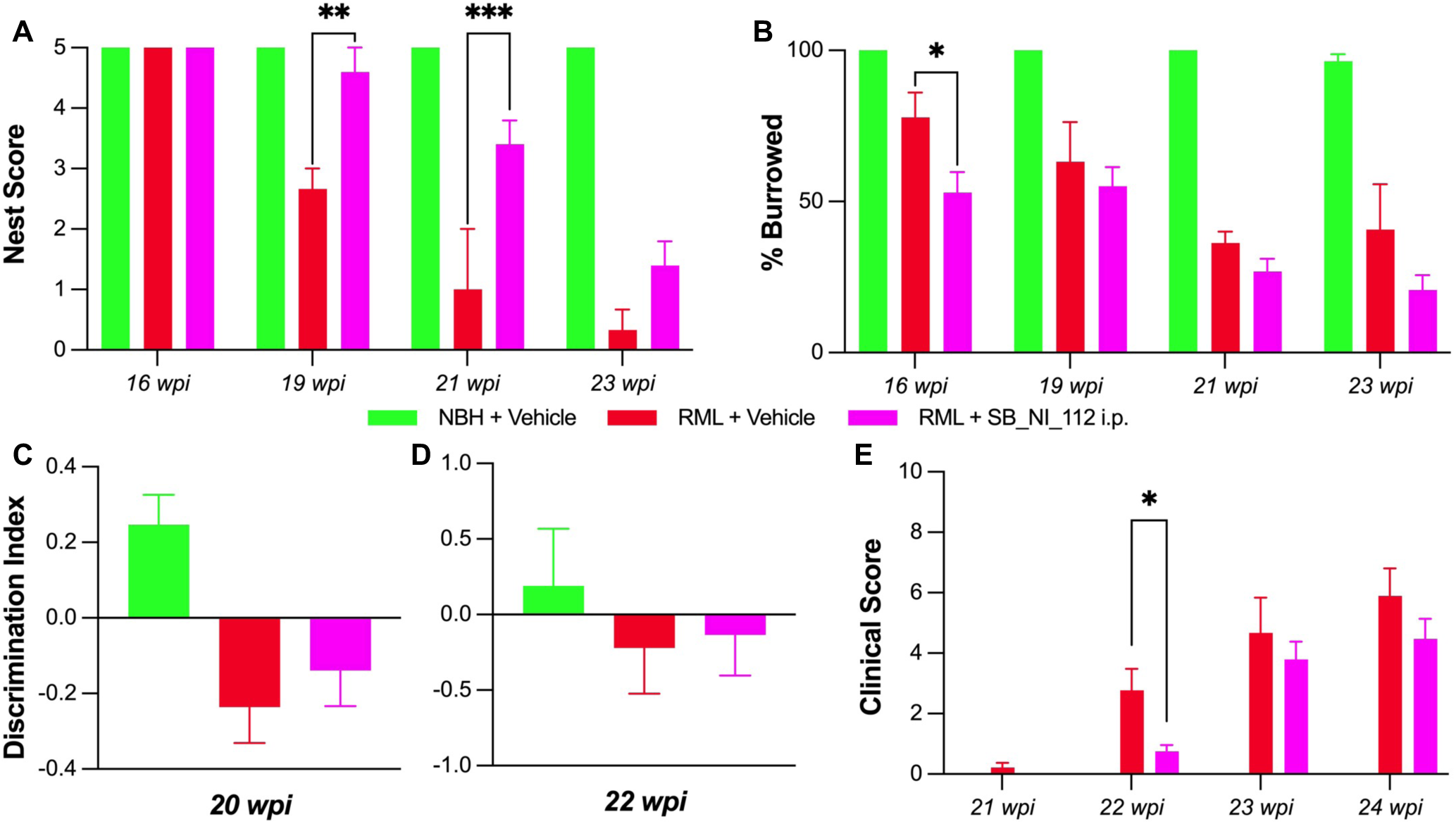
Supporting information contains behavioral and cognitive deficit data analysis at 150 mg/kg 2x/week dosing of SB_NI_112 (Figure S1). Additionally, data for ELISA analysis of IL-28, IL2, IL-1a, and IL-1β (Figure S2). Lastly, videos displaying clinical signs (clasping, righting reflex, and walking) for control and treatment groups at 22 weeks post inoculation (wpi) are available in Supplement Videos 1-9.

## Notes

### Summary of Updates

small edits to text

