## Supplemental file for "Targeting neuroinflammation by pharmacologic down-regulation of inflammatory pathways is neuroprotective in protein misfolding disorders"

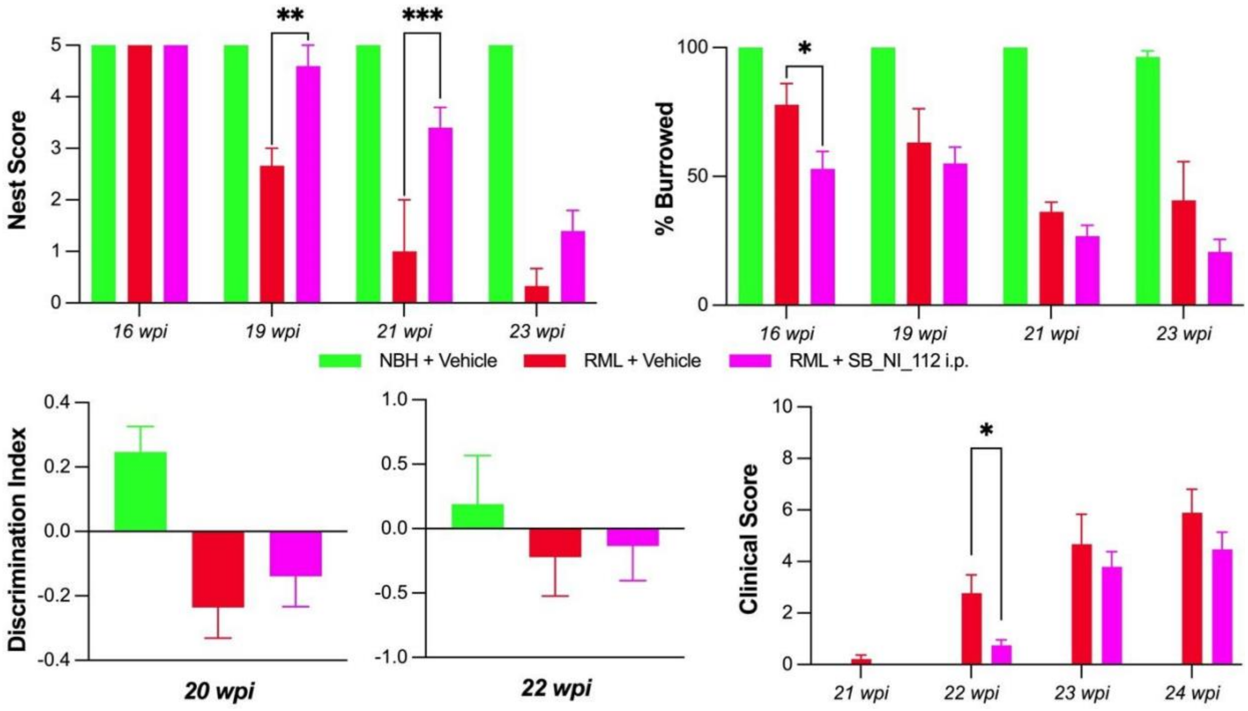


**Figure S1. Behavioral and cognitive deficits are protected by NF-kB and NLRP3 down-regulation with high-dose SB_NI_112 in prion diseased mice.** A significant increase in the ability of the mice to build proper nests at 19 and 20 wpi with high-dose (150 mg/kg 3x per week) SB_NI_112 i.p. compared to vehicle treatments. N=9-16. Two-way ANOVA, error bars = SEM, ** $p < 0.01$, *** $p < 0.001$. In the hippocampal burrowing assay, high-dose SB_NI_112 i.p. treated mice showed significantly better ability to burrow, along with high dose SB_NI_112 i.p. at 16 weeks, compared to untreated counterparts (B). Two-way ANOVA, error bars = SEM, * $p< 0.05$. No significant difference seen in novel object discrimination index at 20 (C) and 22 wpi (D), though trend present towards improvement with SB_NI_112 i.p. N=9-16. One-way ANOVA, error bars = SEM. Clinical scores of mice treated with high-dose SB_NI_112 i.p. at 22 wpi were significantly lower than vehicle treated counterparts with a continued trend towards lower scores through 24 wpi (E).

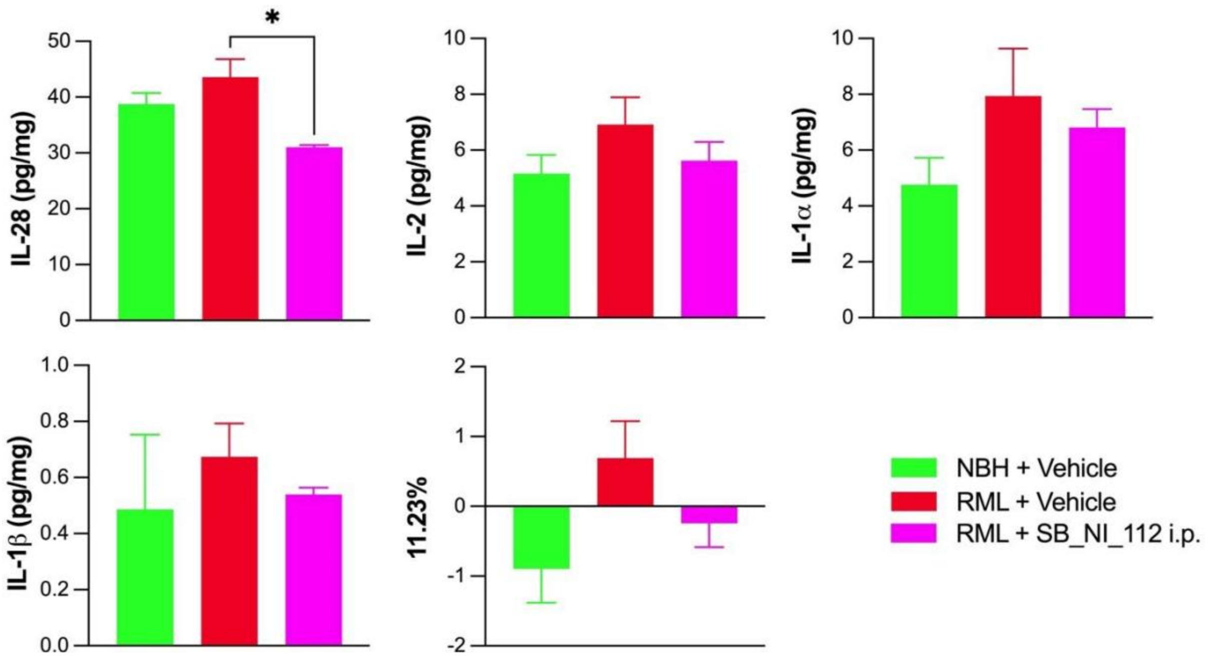


**S2. Inflammatory cytokine profile changes with SB_NI_112 i.p. treatment.** At 16wpi IL-28 (A) is significantly reduced in i.p. SB_NI_112 mice treated mice compared to untreated counterparts. IL-2 (B), IL-1α (C), and IL-1β (D) trend lower, similar to NBH control, with SB_NI_112 i.p. compared to untreated counterparts, measured in pg/mg of total protein. N=4 for all groups, outliers removed with ROUT analysis. One-way ANOVA, error bars = SEM, * $p < 0.05$.

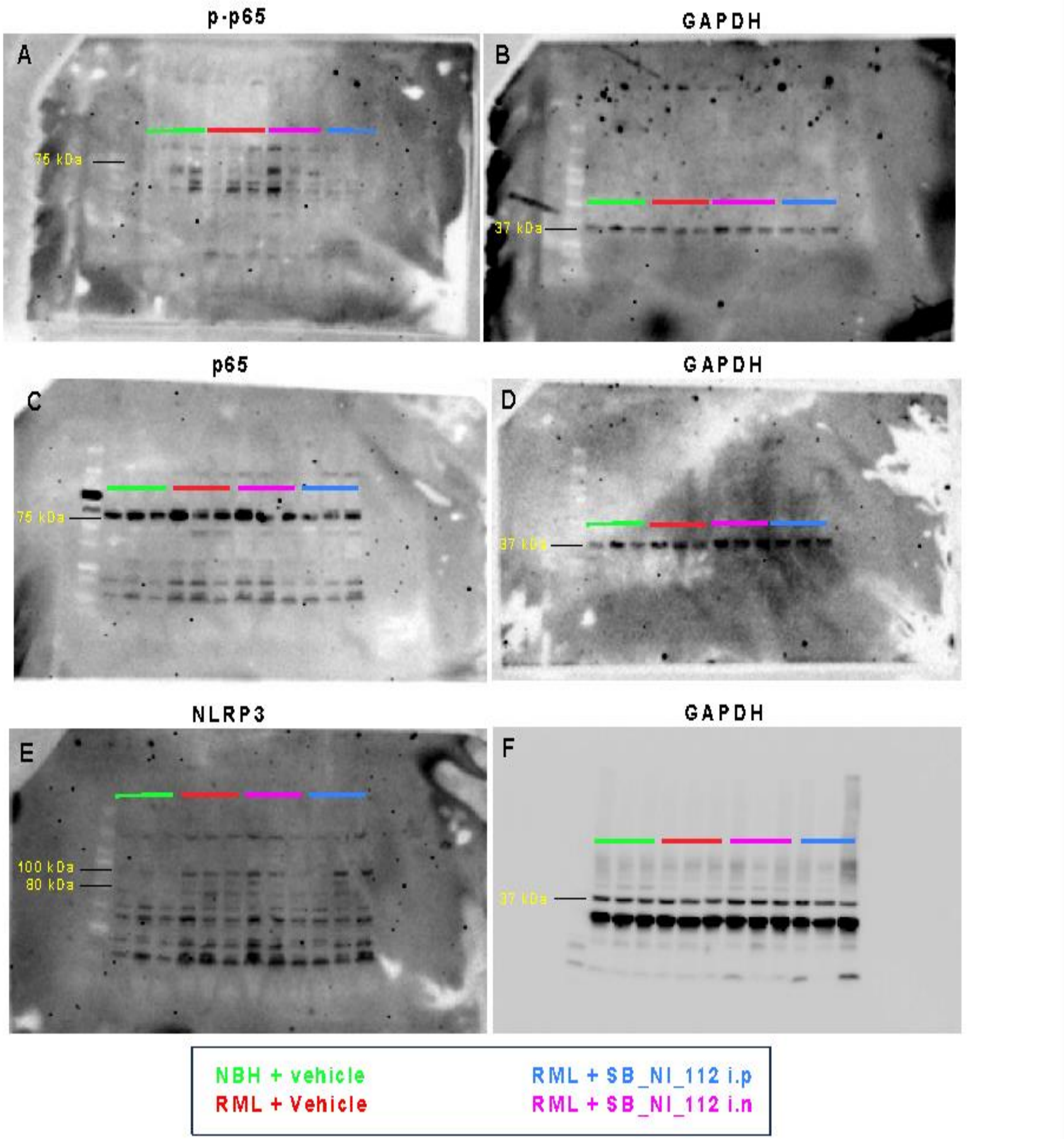


**S3. Original blots for Phospho-p65, p-65 and NLRP3 with SB_NI_112 i.p. and i.n treatment.** At 20wpi brain tissue was immunoblotted for phosphorylation of p65, total p65 and NLRP3 . N=3 for all groups. Uncropped blots used in Figure 3 shown here.

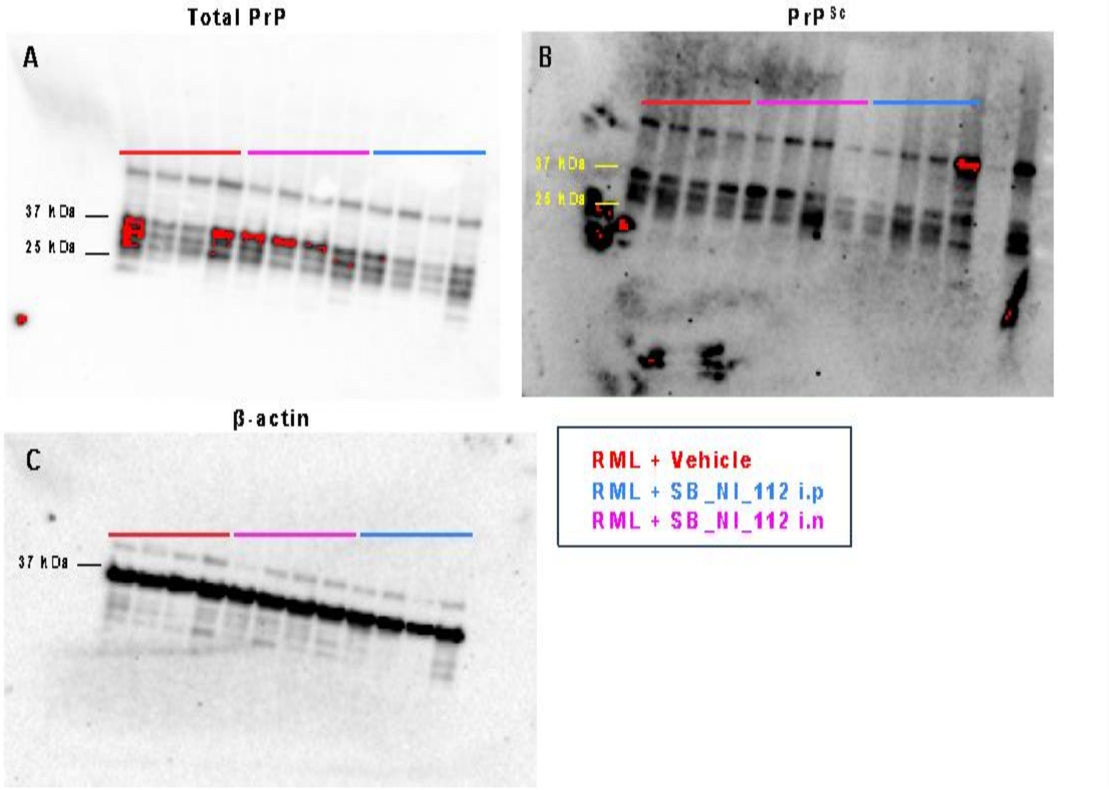


**S4. Original blots for Total PrP and PK resistant material with SB_NI_112 i.p. and i.n treatment.** Terminal brain tissue was immunoblotted for total PrP and PK resistance material . N=4 for all groups. Uncropped blots used in Figure 8 shown here.

**Supporting Videos:** Videos were taken at 22wpi to display clinical signs in control, RML + vehicle, and RML + SB_NI_112 i.p. mice. Videos 1-3 show extensor reflex (clasping), videos 4-6 show righting reflex, and videos 7-9 show mice walking (clinical signs visible through walking include tail rigidity, hyperactivity, ataxia, tremors, kyphosis, and poor grooming).

**Supporting Video 1.** Clasping Control + Vehicle
**Supporting Video 2.** Clasping RML + Vehicle
**Supporting Video 3.** Clasping RML + SB_NI_112 i.p.
**Supporting Video 4.** Righting Control + Vehicle
**Supporting Video 5.** Righting RML + Vehicle
**Supporting Video 6.** Righting RML + SB_NI_112 i.p.
**Supporting Video 7.** Walking Control + Vehicle
**Supporting Video 8.** Walking RML + Vehicle
**Supporting Video 9.** Walking RML + SB_NI_112 i.p.
